# Comparative analysis of the Accelerated Aged seed transcriptome profiles of maize CSSLs (I178 and X178)

**DOI:** 10.1101/627117

**Authors:** Li Li, Feng Wang, Xuhui Li, Yixuan Peng, Hongwei Zhang, Stefan Hey, Guoying Wang, Jianhua Wang, Riliang Gu

## Abstract

Seed longevity is one of the most essential characters of seed quality. Two Chromosome segment substitution lines (CSSL) I178 and X178 with significant difference on seed longevity were subjected to transcriptome sequencing before (0d-AA) and after five days of accelerated ageing (5d-AA) treatments. Compared to the non-accelerated ageing treatment (0d-AA), 286 and 220 differential expressed genes (DEGs) were identified in I178 and X178, respectively Among those, 98 DEGs were detected in both I178 and X178 after 5d-AA, Enriched GO terms included cellular components of cell part, intracellular part, organelle and membrane etc., including carbohydrate derivative catabolic process, carbohydrate synthesis, sugar isomerase (SIS) family protein etc. Transcriptome analysis of I178 and X178 showed that Alternative splicing (AS) occurs in 63.6% of the expressed genes in all samples. Only 381 genes specifically occurred AS in I178 and X178 after 5d-AA, mostly enriched in nucleotide and nucleoside binding. Combined with the reported QTL mapping result, the DEG and the AS information, 13 DEGs in the mapping intervals and 7 AS-DEGs were potential candidates may directly or indirectly associated to seed ageing.

## 1. Introduction

Ageing is an inevitable process affecting seed longevity, the length of time a seed remains viable, which is accompanied with a progressive loss of quality or viability over time, and a crucial issue for germplasm conservation and seed marketing [1]. Seed longevity depends greatly on seed moisture, relative humidity, oxygen pressure and temperature of storage [2, 3]. Seed storability is a complex trait, many studies have illustrated the oxidative and mitochondrial damage occurs during seed storage, which are the reasons for loss of longevity, and. During seed storage, the activity of the ascorbate and glutathione (AsA-GSH) cycle is reduced resulting in ROS accumulation [4, 5], and carbonylation happened of the proteins in seeds [6, 7]. These play important roles in scavenging reactive oxygen species (ROS). High concentration of H_2_O_2_ and the malondialdehyde (MDA), end-products of lipid peroxidation, were considered as the critical factor in contributing to seed deterioration and influencing seed longevity and vigor [8–11]. Moreover, the activity of mitochondria, the organelles that supply energy for seed germination, is significantly decreased during storage, thereby inhibiting seed germination [5, 12]. Thus, the level of antioxidants and the level of energy supply are key factors in the regulation of seed longevity. The genetic architecture of crop seed longevity is still far behind the model plant Arabidopsis. Recently, various omic approaches have been used to investigate protein expression patterns during seed storage [13, 3, 14, 15, 7, 16]. Sano et al performed an RNA-seq experiment on bulked RILs of Est-1 × Col-0 in Arabidopsis and found that brassinosteroid is important for seed longevity by regulating the seed coat permeability by BR signaling pathway [13]. Lv et al conducted proteomic analysis on artificially aged wheat seeds; differentially expressed proteins (DEPs) were mainly involved in metabolism, energy supply, and defense/stress responses. Up-regulated proteins were mainly enriched in ribosome, whereas the down-regulated proteins were mainly accumulated in energy supply, starch and sucrose metabolism and stress defense (ascorbate and aldarate metabolism), revealed that the inability to protect against ageing leads to the incremental decomposition of the stored substance, impairment of metabolism and energy supply, and ultimately resulted in seed deterioration [14]. Yin et al performed proteomic analysis on rice embryo with different days of ageing, by comparison analysis, they found most of down regulated proteins were related to energy metabolism (29%), defense (21%), glycolytic pathway (8%), protein synthesis (8%), protein destination and storage (6%), transcription (5%), growth or division (4%), secondary metabolism (3%), transporting (1%), signal transduction (1%). While most of the upregulated proteins were related to storage, energy, disease and defense, metabolism, protein synthesis, growth and division [7]. Chen et al compared the storage ability of two wheat varieties (storage tolerant vs. storage sensitive), most of the proteins remarkably different from two varieties were mainly associated with disease or defense, protein destination and storage, energy, also the storage tolerant seeds possessed a stronger ability in activating the defense system against oxidative damage, utilizing storage proteins for germination, and maintaining energy metabolism for ATP supply [16]. While seed longevity is different between species, it also differs between genotypes of a species, the genetic information that involving in seed longevity was far beyond the understood. Also the expression of genes and proteins related to disease defense and energy is largely altered during seed storage. However, the effect of such changes on ageing tolerance and sensitivity of seeds is largely unknown.

In the present study, the simulation of natural seed deterioration by artificial accelerated ageing (AA) treatment by controlled the moisture up to 95% and the relative temperature at 45 °C. Two chromosome segment substitution lines (CSSL) (X178 and improved 178, I178), with similar genetic background but shows different sensitivity or endurance in terms of ageing process, were subjected the transcriptional expression analysis, differential expressed genes (DEGs) of non-accelerated ageing treated dry seeds (0d-AA) and 5 days accelerated ageing treatment (5d-AA) in two 178 lines were detected and the co-DEGs as well as the genotype-specific DEGs were subjected qRT-PCR validation, followed by alternative splicing analysis to discover genes or pathways that affected during seeds ageing. We were aiming for uncovering ageing related genes by DEG, alternative splicing (AS) and the candidate gene analysis which existed in QTL mapping interval. Consequently, exploring more biology process that been affected by ageing or those processes that influencing the ageing.

## 2. Materials and Methods

Maternal parent X178 was a widely cultivated maize hybrid Nongda108 (released by China Agricultural University in 2001), an elite line which has better agronomic trait of storability. I178 (Improved X178) was derive from X178 by introgression of chromosome segments for several generations (CSSL), followed by consecutive self-crossing for at least 10 generations.

### Accelerated ageing treatment (AA)

The simultaneously fresh harvested (FH) I178 and X178 seeds were surface disinfected with 1% NaClO for 5 min and washed 10 times with sterile-distilled water, balancing the moisture in the room temperature for overnight, followed by accelerated aging treatment of suspending the seeds on metal mesh trays within closed metal boxes (25× 25×14 cm), maintained in an ageing chamber (LHC-150-11, Beijing Luxi Ltd) at the condition of 95% moisture content and 50 °C temperature for 3, 5 and 7 days of different purpose, 3 replications and non-accelerated aged (0d-AA) seeds as the control. The storage condition of harvested seeds was under 10 °C in a cold room.

### Protein Quantification and Zein analysis

Samples of I178 and X178 were prepared for zein extraction, including: 1). dry seeds; 2). seeds after 6 hours imbibed water (0d-6h); 3). Seeds germinated 48 hours (0d-48h); 4). seeds after 3d-AA; 5). Seeds after 5d-AA; 6). seeds after 7d-AA were prepared as following methods: dry seeds without any treatment, imbibed into water for 6 hours, 48 hours seeds germination was performed with pre-wetted crepe cellulose papers (CCP), and covered with another piece of CCP, rolling into paper rolls and upright in the ziplock bags for 48 hours under 25 °C. Above samples were powdered in liquid N_2_ and 50 mg of powder was used for zein extraction. Removing the lipids with petroleum benzin and dissolving the samples with protein extraction buffer (12.5 mM sodium borate, 2% 2-mercaptoethanol, 1% SDS and pH10), followed by 5 min incubation at 37 °C, centrifuged at 14,800 rpm for 15 minutes, the supernatant contains total protein was incubated with absolute ethyl alcohol for 2 hours. After centrifugation at 14,800 rpm for 15 minutes, the supernatant contains zein was dissolved in IPG solution (8 M urea, 220 mM DTT and 2% CHAPS) and measured with the BCA protein assay kit (TRANS, Beijing).

### Tetrazolium chloride (TTC) staining

By refer to the Tetrazolium staining method (TZ) on soybean (*Glycine max*.) vigor test [48], corn seeds were imbibed with water for 20 hours at room temperature prior to staining, cut the seeds longitudinally through embryo, then staining with 0.1% Tetrazolium chloride solution (TTC, aqueous solution of 2,3,5-triphenyl tetrazolium chloride) for 1 h and washing 3 times before observation.

### RNA-seq and qRT-PCR

For RNA-Seq experiment, 100 artificial accelerated aged seeds (5d-AA) were pooled and grinding promptly in liquid nitrogen, 0.1 g of powder was used for isolating the mRNA with the RNAprep pure Plant Kit (Cat#DP432, TIANGEN, Beijing), RNA was quality check the total RNA with the 2% agrose gel, high quality RNA was used for RNA-Seq library preparation and sequenced on a Illumina HiSeq2500 platform (Berry Genmics, Beijing). Two biological replications included and the Non-accelerated aged dry seeds (0d-AA) as the control.

RNA for qRT-PCR experiment was extracted as above procedure, Quality checking the RNA and performed the reverse transcription with the OneScript cDNA Synthesis Kit (Cat#G234, ABM, Canada), primer of the genes was designed with software Primer Premier5.0. The Fast Sybr Green Master Mix (Applied Biosystems, Foster City, CA, USA) was employed, according to the manufacturer's instructions, in a reaction volume of 10 μl. qRT-PCR was conducted on a ABI Quantstudio™ DX Real-Time PCR system (Applied Biosystems). PCR conditions included initial denaturation for 2 min at 95 °C, followed by 40 cycles of denaturation at 95 °C for 30 s, hybridization at 60 °C for 40 s, and elongation at 68 °C for 10 s. The actin2 gene was used as an internal control. The 2^−ΔΔct^ method was used to calculate the relative level of gene expression, and the B73 sample served as a control. A relative level of gene expression greater than 1 was considered to indicate up-regulation, and less than 1 indicated down-regulation. All qRT-PCR reactions were performed with the three biological replicates.

### Data analysis

For gene expression level in I178 and X178, transcription with FPKM (Fragments per Kilo bases per Million fragments mapped) >0.1 was considered as expressed genes calculated by htseq-count in HTSeq software. To identify genes involving in seeds ageing, the comparison of genes expressed after 0d-AA and 5d-AA was performed in both I178 and X178, DESeq2 was used for differential expression analysis with the Fold Change of 1.5, with adjusted P-value (*q-value)* <0.05 as the threshold value. Venn diagram was performed with online software (http://bioinfogp.cnb.csic.es/tools/venny/index.html) [49]. Agri-Go enrichment was also performed with online (Agri GO v2.0; http://systemsbiology.cau.edu.cn/agriGOv2/#) database [50]. Gene Organ or tissue specific expression level was compared with online q-teller database ((http://www.qteller.com/qteller4/)).

## 3. Results

### 3.1 Comparison of seeds storability for I178 and X178

Few morphological and the physical differences were observed between X178 and I178 because of the similar genetic background. Seed storability can be reflected by color of seed coat, seed viability and vigor after long term storage or AA treatment. Seed coat of I178 was obviously oxidized after 5d-AA as the brown color, and the seed viability was reduced dramatically in I178 based on triphenyl tetrazolium chloride (TTC) staining, fresh harvested (FH) seeds have highest viability and dehydrogenase activity as the embryo part stained with bright-color while light-colored after 3d-AA and especially no color after 5d-AA (Fig 1A, B). FH seeds of two lines showed slight difference of relative conductivity (RC) before 3d-AA, significant difference observed after 5d-AA. After 1-year storage, the RC of I178 was two times higher than X178, as the continued AA treatment, the storability was reduced significantly in I178 after 5d-AA (Fig 1C). Comparative SDS-PAGE analysis of seed storage protein (SSP) zein in dry seeds, imbibed seeds (6 hours imbibition), germinated seeds (48 hours germination) and 3, 5 and 7 days accelerated aged seeds (3d-AA, 5d-AA and 7d-AA), and no significant difference was observed between two 178 lines after AA treatment except the degradation of 40 kD protein happened in I178 after 5d-AA, after 7d-AA, the proteins smaller than 25 kD (including γ27, α22, α19, γ16, β15 and δ10) were also dramatically degraded in I178 (Fig 1D).

**Fig 1.**
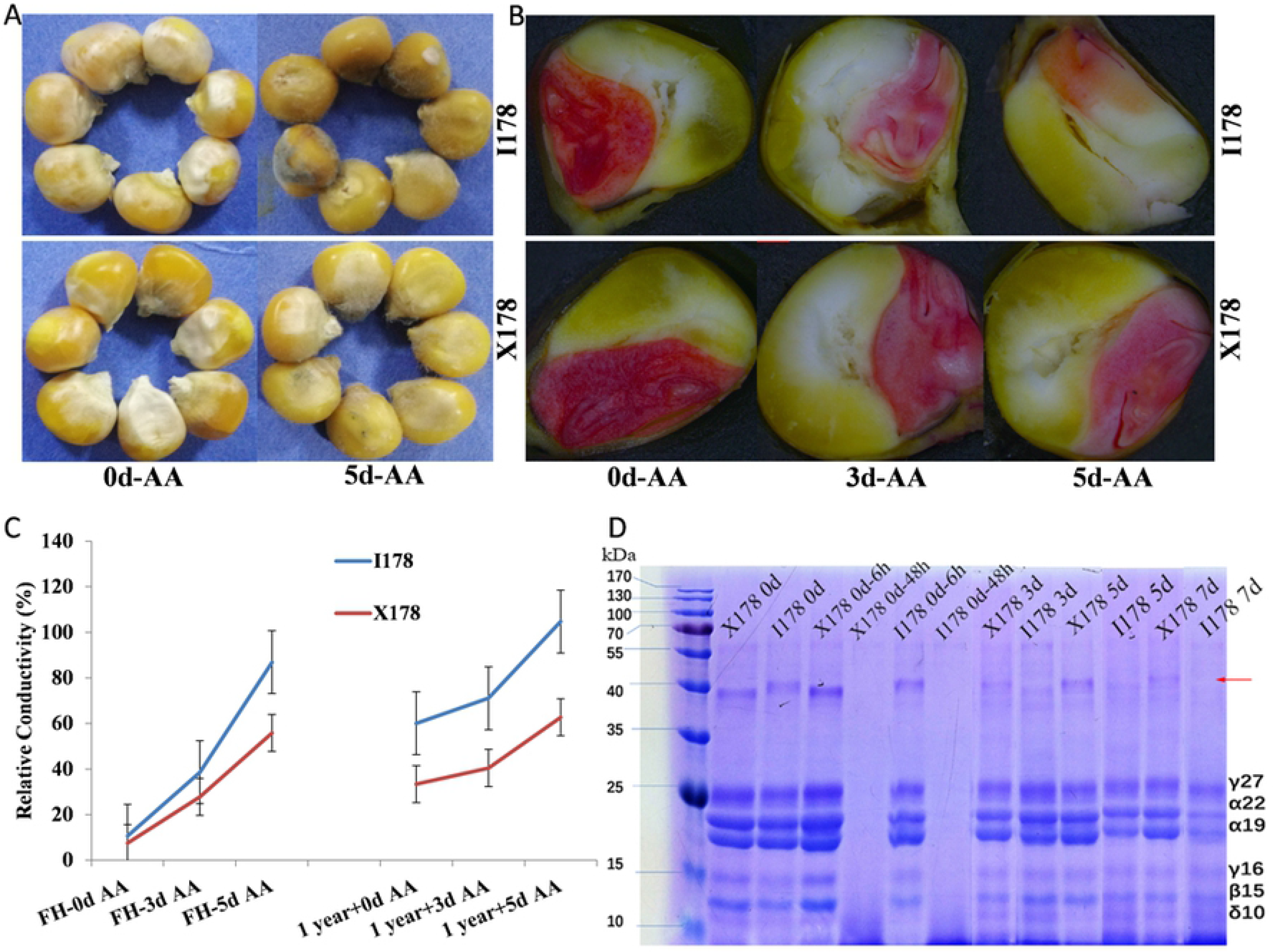
Storage characterization of I178 and X178 seed. **A).** Observation of 3 and 5d-AA seeds; **B).** TTC staining of embryo at 0d-AA, after 3 and 5d-AA; **C).** The relative conductivity of fresh harvest (FH) seeds, 1-year storage seeds treated by 3 and 5d-AA; the dash lines indicate the vigor of FH seeds after 5d-AA was lost more than that of 1-year storage seeds. **D).** SDS-PAGE analysis of seeds zein, lane 1-12: Dry seeds of X178 and I178 (X178-Dry, I178-Dry), X178 seeds after 6 hours imbibed water (X178-0d-6h), 48-hours germinated X178 seeds (X178-0d-48h), I178 seeds after 6 hours imbibed water (I178-0d-6h), 48 hours-germinated I178 seeds (I178-0d-48h), X178 seeds after 3d-AA (X178-3d), I178 seeds after 3d-AA (I178-3d), X178 seeds after 5d-AA (X178-5d), I178 seeds after 5d-AA (I178-5d), X178 seeds after 7d-AA (X178-7d), I178 seeds after 7d-AA (I178-7d). The degradation of 40 kD protein was denoted as arrow.

### 3.2 Transcriptome profile of I178 and X178 seeds

The cDNA libraries of the non-accelerated aged dry seeds (0d-AA), 5 days of accelerated aged seeds (5d-AA) for I178 and X178 were prepared and sequenced using an Illumina HiSeq 2500 platform, 8 Gb data of two biological replicates for each sample were obtained. Reads of low sequencing quality were filtered out (about 0.12%~0.18%), and totally 34.4~43.4 million 100 bp paired-end reads were generated in I178 and X178, with the average of 40.8 million reads in each sample. At least 97.22% high quality reads used for analysis. Of these reads, average 68.8% unique mapped reads (289.9 million) were aligned to the B73 reference genome to estimate the transcript levels (ZmB73_RefGen_v3). Expression values were calculated with units of fragments per kilo-base per million reads mapped (FPKM). As expected, 92.92% high quality reads can be mapped to protein-coding genes, 1.36% and 5.72% were mapped to intron and intergenic region, respectively **(Table S1)**.

### 3.3 Identification of the DEGs in I178 and X178

Genes with same expression patterns during seed ageing may related to ageing metabolic processes [17]. To identify the function of genes, researchers clustered genes with similar expression patterns as clues to study the function of unknown genes [18]. The RNA-Seq reads of 8 samples for I178 and X178 after 0d-AA and 5d-AA (2 replications) were aligned to the maize reference genome (ZmB73_RefGen_v3), the reads coverage and the correlation between 2 replications was relatively high which reflected by the FPKM distribution and pearson correlation as shown in supplementary **(Fig S1)**. The expressed genes in I178 and X178 were 26,909 and 26,514, respectively, among that 25,242 common genes (more than 94%) expressed in both 178 seeds (FPKM > 0.1). To identify genes involving in seeds ageing, we focused on genes that differential expressed after 5d-AA (compared to 0d-AA), totally 286 and 220 DEGs were detected in I178 and X178, and 98 common DEGs in both I178 and X178 (log2FC ≥ 0.585). Only 2 common up-regulated and 86 down-regulated genes identified in two 178 (Fig 2C; **Table S2**).

**Fig 2.**
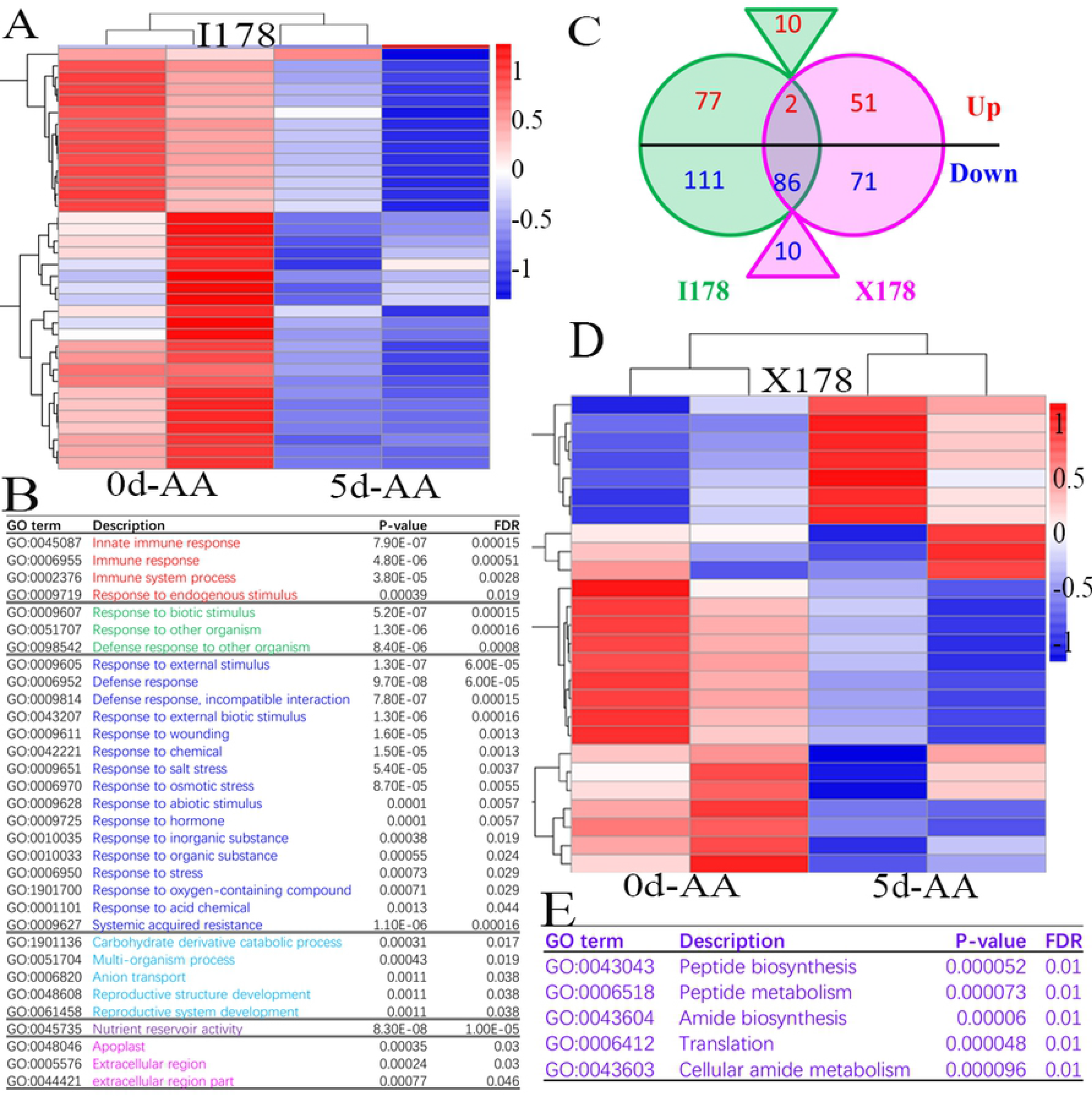
RNA-Seq of I178 and X178 and gene expression analysis before and after 5d-AA. **A).** Heatmap of differential expressed genes in I178 after 5d-AA compared with 0d-AA. **B).** Most of the significantly differential expressed genes in I178 were enriched in 5 categories of Immune system (red), biotic stress response (green), abiotic stress response (blue), carbohydrate catabolic process (light blue), nutrient reservoir activity (purple) and extracellular region (pink). **C).** Compared to the 0d-AA treatment, DEGs after 5d-AA identified in two materials with the adjusted *P*-value ≤0.05 and the FoldChange ≥1.5, there are 188 specific I178 DEGs and 122 X178 specific DEGs, among that, 98 common DEGs identified in I178 and X178, with 2 common up-regulated and 86 common down-regulated genes. 10 genes were up-regulated in I178 while down-regulated in X178 was labeled in the triangle. **D).** Heat map of DEGs in X178. **E).** GO enrichment of the most significantly differential expressed genes in X178.

### 3.4 GO enrichment of DEGs after 5d-AA

GO is an internationally standardized gene function classification system used to describe the properties of genes and their products in any organism, which contains three ontologies: biological process, cellular component and molecular function [19]. The 286 I178 differential expressed genes were mainly involving in biology process of abiotic response like temperature, salt stress, osmotic stress, light intensity, heat, ethanol, abiotic stimulus, heat acclimation etc., and the molecular function of nutrient reservoir activity, chitin binding and catalytic activity **(Fig S2A).** While for the 220 X178 DEGs were mainly enriched in cellular component of nuclear part, organelle lumen, intracellular part and heterochromatin etc. (**Fig S2B**).

Most of the down-regulated genes in I178 were enriched in stimulus response, including response to biotic stress (organism stress, external biotic etc), abiotic stress (inorganic substance stress, chemical, oxygen-containing compound, acid chemical, organic substance, endogenous stimulus, hormone etc), immune response (defense response, innate immune etc.), and most of DEGs in X178 were enriched in DNA or protein biosynthesis process like peptide biosynthesis, amide biosynthesis and translation (Fig 2A-B; D-E). The common 98 DEGs in two 178 were mainly enriched in cellular component of cell part (GO:0005623; GO:0044464), membrane-enclosed lumen (GO:0031974), organelle and intracellular organelle (GO:0043226; GO:0044422; GO:0043233; GO:0043229) and intracellular part (GO:0044424; GO:0070013; GO:0044446; GO:0043231), RNA polymerase (GO:0030880; GO:0000428; GO:0055029; GO:0016591), membrane bounded organelle (GO:0043227), transferase complex (GO:0061695) and the nuclear part (GO:0005634; GO:0044428; GO:000319981; GO:0005654), most of DEGs were down-regulated, and enriched in biology process of carbohydrate derivative catabolic and molecular function of carbohydrate derivative binding (**Fig S2D, E**). Only two common up-regulated genes in I178 and X178, gene GRMZM2G353885, encodes a TATA box binding protein (TBP) associated factor 2, and another gene of no annotation (**Fig S2C**).

### 3.5 Analysis of DEGs by qRT-PCR

According to the RNA-Seq results, 9 genes including 7 randomly selected DEGs and 2 longevity related genes were selected for qRT-PCR validation. The expression patterns of the 7 DEGs were consistent between qRT-PCR and RNA-seq, indicating that the RNA-seq gene expression was reliable, two genes (ZmLOX11 and ZmPIMT1) were not able to detected in RNA-Seq analysis, further qRT-PCR was conducted on I178 and X178 before and after 5d-AA treatment, as expectation, the expression was extremely low, and was consistent to the q-Teller whole transcriptome expression result (http://www.qteller.com/qteller4/) (Fig 3).

**Fig 3.**
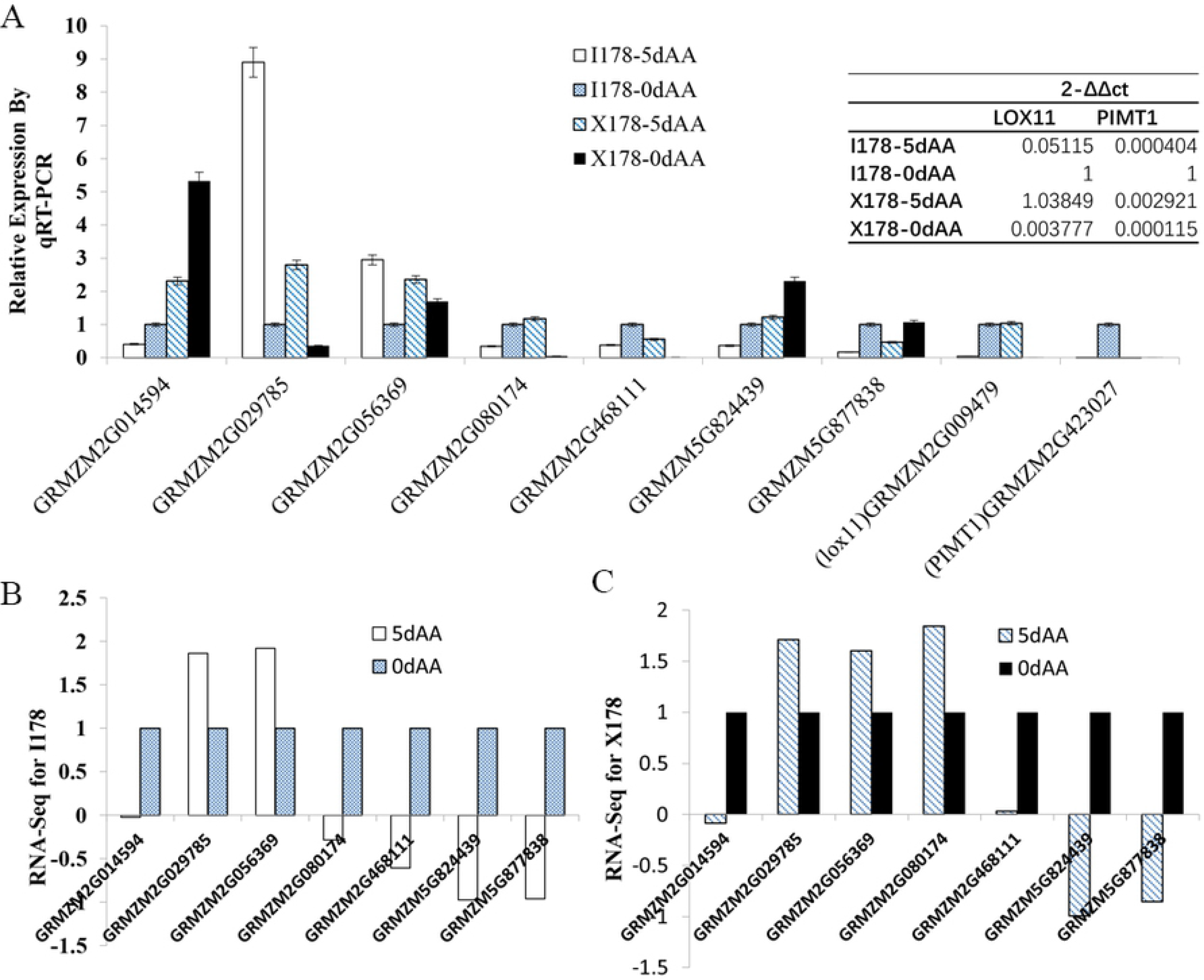
qRT-PCR validation of the RNA-Seq result. **A).** Nine genes including 7 differential expressed genes and 2 ageing related genes validated by qRT-PCR. RNA-Seq gene expression calculated by fold-change in I178 **(B)** and X178 **(C)**.

### 3.6 Alternative splicing (AS) analysis of ageing related transcriptions and GO enrichment

In I178 and X178, we totally identified 51,388~59,146 AS events, including 12 different types which covered 46,521~ 48,724 transcript isoforms (including 15,984~17,070 genes) **(Table S3)**. Among that, TSS (alternative 5’ first exon) and TTS (alternative 3’ last exon) account for more than 72% of the total AS events, and IR (Intron retention), AE (Alternative exon) and SKIP (Skipped exon) were also frequently occurred in I178 and X178 (**Fig S3A**). In dry seeds (0d-AA), we detected 56,228 and 53,593 AS events in I178 and X178, respectively, which cover 20,446 and 19,623 transcripts in I178 and X178, respectively. After 5d-AA, we detected 55,763 and 56,794 AS events in I178 and X178, which cover 20,073 and 19,845 transcripts, respectively **(Fig S3A; Table S3)**. By comparing AS in I178 and X178 before and after 5d-AA, we noticed that 63.6% transcript isoforms (15,606, cover 12,834 genes) occurred AS in all samples. In order to discover that AS genes may involving in seed ageing, we only focus on AS genes that specifically identified in I178 and X178 after 5d-AA: 381 transcript isoforms (including 169 genes) occurred AS in both 178 lines, 849 transcript isoforms (including 415 genes) specifically occurred AS in I178 and 760 transcript isoforms (including 343 genes) specifically occurred AS in X178 after 5d-AA (**Fig S3B**). In order to identify the relationship of AS genes and the ageing related procedure, Agri-Go analysis on common AS genes in I178 and X178 showed that the enriched in nucleotide biosynthesis process, function as nucleoside binding (Fig 4A). For those AS genes specifically occurred in I178 and X178 after 5d-AA, beside of the molecular function of ribonucleoside binding, ATP binding etc., some were also enriched in freezing response (Fig 4B). Combine the DEG and the AS gene information in this study, only 6 X178 specific DEGs specifically occurred AS and one I178 specific DEG specifically occurred AS after 5d-AA (Table 1; **Fig S3C**).

**Table 1.**
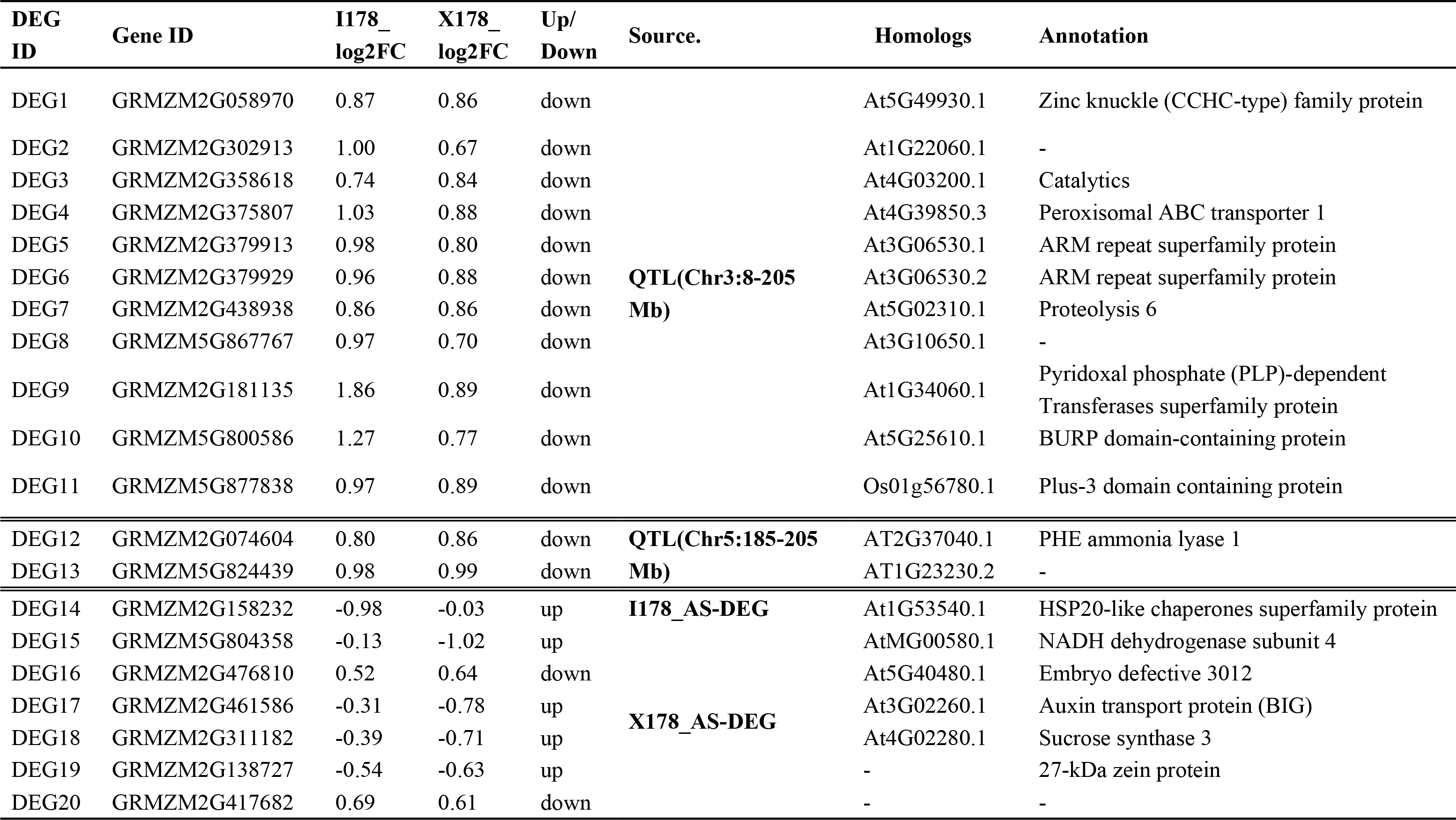
Potential seed ageing related genes.

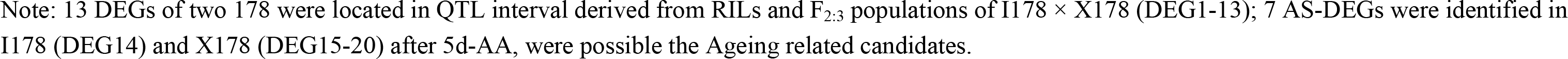

**Fig 4.**
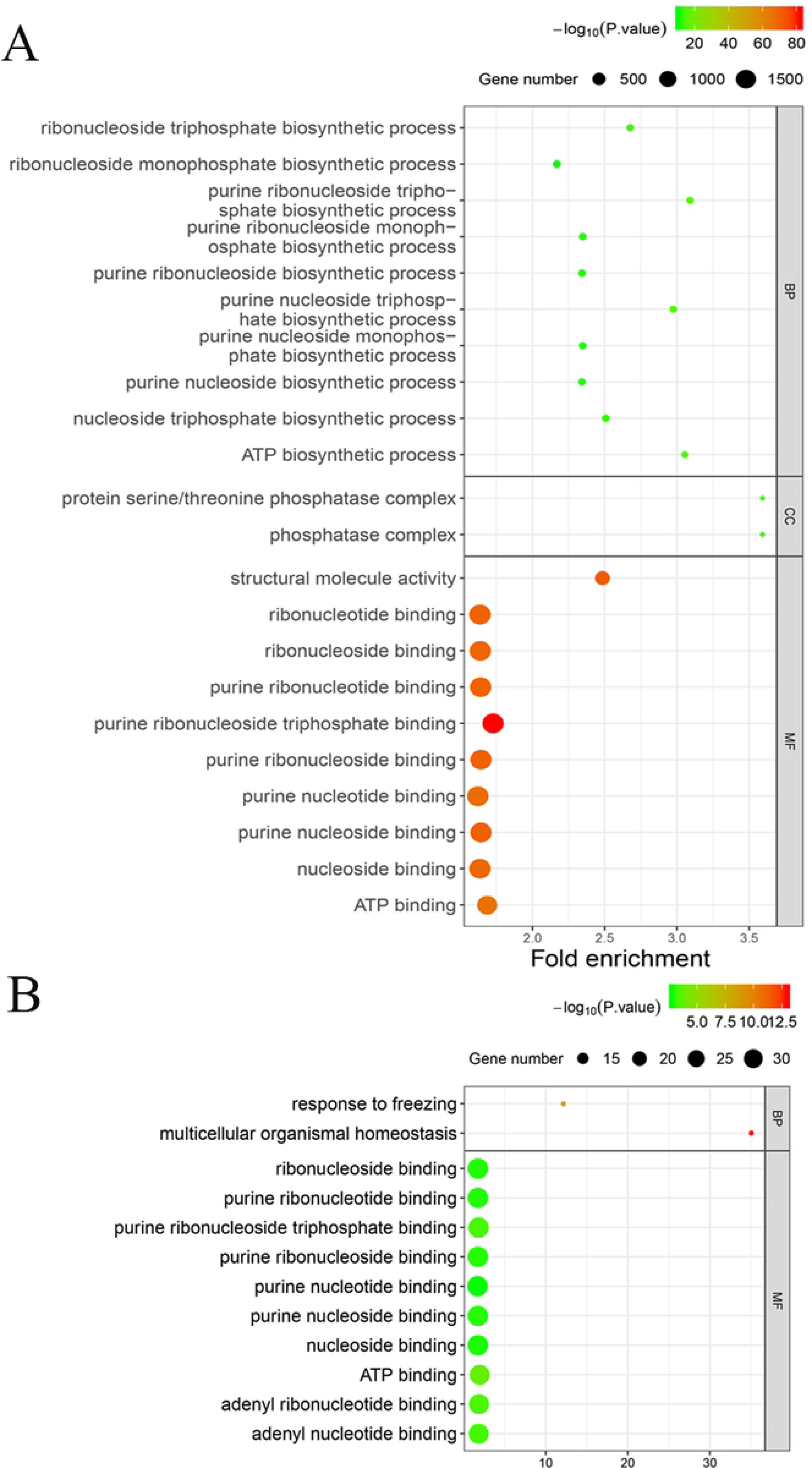
GO enrichment of AA induced alternative splicing genes (ASG) in I178 and X178. **A).** Biology process, cellular component and the molecular function of common ASGs in I178 and X178. **B).** 381 alternative splicing genes specifically expressed after 5d-AA in I178 and X178.

## 4. Discussion

### 4.1 Seed storability was decreased dramatically in I178 after AA treatment

I178 was derived from X178 and less polymorphic segments was detected among the chromosomes except chromosomes 7 and 10 by performing 6K maize array chip, the only significant difference between X178 and I178 is the seed vigor which was validated by previous study [20]. The electrolytic exudate conductivity of the seed reflects the permeability of the seeds coat, also represent the damage of cellular membrane system, our test of seed vigor of FH seeds after 3d-AA and 5d-AA was consistent to Liu’s result. Obviously, the seeds storability was constantly decreased even stored at relative low temperature, and the ageing level of fresh harvested seeds after 5d-AA was stronger than 1-year storage in cold room (Fig 1). As we know the deterioration exists inevitably, to slow down the loss of longevity, keep the seeds in a lower temperature (possible 4 °C) is a better choice. Seeds storage protein (SSP) have been described as a primary target for oxidation in seeds, zein is the largest component of SSP which account for 60% of SSP [21]. Carbonylation of SSPs during long-term storage was an irreversible form of oxidation leading to deterioration in both after-ripen seeds and aged seeds [22, 23]. More studies found that SSPs were degraded during seed germination, which also differential expressed in aged seeds, indicating the role of SSPs in seed longevity [24, 6, 7, 16]. In our study, we do observed the degradation of the SSPs (zein) after 48 hours of germination in both I178 and X178. Previous study proved that the oxidative SSPs were more easily degraded into smaller polypeptides or amino acids [16], in our observation, the total amount of zein was decreased significantly after 5d-AA in I178 and 7d-AA in both 178 lines, was consistent to above result.

### 4.2 Seed ageing affects numerous biology processes including stress defense and carbohydrates metabolisms

Seed ageing is an inevitable procedure occurred on all living things. The consensus molecular mechanism associated to the ageing is including: 1) Peroxidation of plasma membrane and disintegration of membrane system structures. 2) Variation of biomacromolecule, including variation of nucleic acid (RNA and DNA) and enzyme/proteins. 3) The accumulation of toxic substances, i.e., Reactive oxygen species (ROS), malondialdehyde (MDA), and the by-products of seeds physiological activity, organic substance like alcohols, free fatty acids etc. [25, 16]. In our RNA-Seq analysis 5d-AA treated 178 lines, 98 common DEGs were identified in both 178 seeds, while only 88 genes have same expression pattern in I178 and X178, 86 of which were down-regulated and enriched in carbohydrate derivative catabolic process **(Fig S2E)**. Interestingly, Lv et al revealed that the proteins in carbohydrate derivative biological pathway were up-regulated in aged wheat seed especially the up-regulation of genes in amide biosynthesis pathway, and the genes in defense- and stress response were down regulated after ageing [14]. In our study, most of the DEGs (including up- and down-regulated genes) were enriched in defense- and stress pathways in I178 after ageing. The inconsistency between two studies can be explained by the different ageing level of the wheat seeds and maize seeds, in Lv’s study, the germination rate of aged wheat was lower than 20% while based on Liu’s result, the GR of 5d-AA treated I178 and X178 was around 20% and 80%, respectively [20]. We do observed a enrichment of up-regulated genes in carbohydrate catabolic process in I178 (GO:0016052), while down-regulated in X178, i.e., GRMZM2G176307, which encodes a glyceraldehyde-3-phosphate dehydrogenase C2. In Xin’s proteome analysis on maize, they mentioned that the carbohydrates utilization was important in seed ageing and seed vigor, was been influenced in aged seeds [4], which was also consistent to our results. The X178 down-regulated genes were enriched in amide and peptide biosynthesis after 5d-AA (Fig 2E), which was inconsistent with Lv’s result, it was possible that X178 possess a better resistant to ageing, after 5d-AA treatment of, the DNA and protein repair systems was slightly affected which reflected by the down-regulation of ageing affected genes, while as the constant ageing treatment once GR was below 20%, some of the stress response genes will be activated and up-regulated to coping with DNA and protein damage.

### 4.3 ZmPIMT1 and LOX11 were down-regulated after 5d-AA

To validate the RNAseq data, qRT-PCR was performed on 9 of randomly selected genes may related to ageing and consistent results were obtained on both platforms. Protein-L-isoaspartyl (d-aspartyl) O-methyltransferase (PIMT), a typical protein repair methyltransferase related to seed longevity by recognizes isoAsp residues in proteins or peptides and catalyzes the transfer of a methyl group from S-adenosyl methionine (AdoMet) to the free a-carboxyl group of abnormal L-isoAsp residues (as well as the b-carboxyl group of D-aspartyl residues) [26, 27]. There are two PIMT orthologues in Arabidopsis and maize, in this study, ZmPIMT2 share 65% sequence identity with the AtPIMT1, was not affected in both I178 and X178, ZmPIMT1 share 71% sequence identity to AtPIMT1, was not been detected in RNA-Seq experiment. qRT-PCR of PIMT1 showed that ZmPIMT1 was almost no expression signal in 178 seeds in all samples (Fig 3), indicating that the spatio-temporal expression specificity of PIMT1 in dicotyledon and. Monocotyledon plants. Lipoxygenase (LOX) was also a typical longevity related protein which associated to the lipid oxidation in seed or other tissue [28]. There are 13 ZmLOX gene have been identified in maize so far, but few of them have been cloned or further studies in molecular biology level [27]. In Arabidopsis. LOX2 is essential for formation of green leaf volatiles and five-carbon volatiles [29], the homolog ZmLOX11 in this study was no expression in RNA-Seq as well, which is normal since the gene expression in q-Teller showed that this gene was only highly expressed in the young seeds while little expression observed in mature seed **(Fig S4)**. qRT-PCR showed that ZmLOX11 was down-regulated in I178, while up-regulated in X178 after 5d-AA treatment, a possibility that LOX11 was not specific expressed in seeds, or the spatiotemporal specificity of the LOX11 in tissue except the seeds (Fig 3).

### 4.4 Identify genes potentially associated to seed longevity

Numerous studies reported that the seed longevity genes may involve in switching off metabolic activity in seeds, repair systems during seed imbibition and DNA, RNA or protein repair systems [25]. In previous study, QTL mapping of seeds ageing traits on RILs and F_2:3_ populations of I178 × X178, 17 QTL were identified on 5 chromosomes [20]. In order to excavating genes that involving in seed longevity, DEGs in QTL mapping intervals for both I178 and X178 were selected for analysis, 13 DEGs located in the mapped QTL of chromosome 3 (11 genes) and chromosome 5 (2 genes), for the 10 DEGs with explicated annotations, DEG4 encodes a peroxisomal ABC transporter 1, previous study showed that the peroxisomal ABC transporter in plant was essential for transporting hydrophobic fatty acids and large cofactor molecules (carrier for ATP, NAD and CoA), and play an indispensable role in pathways like fatty acid β-oxidation, photorespiration, and degradation of reactive oxygen species [30], it was possible that during seed ageing, the accumulation of reactive oxygen species in seed resulted the down-regulation of DEG4. DEG7 encodes a proteases 6, In Arabidopsis, the aspartic protease 1 (ASPG1) was affected the seed longevity and germination by the process of proteolysis [31], proteases 6 was the major cellular machinery of proteolysis in eukaryotic organisms, it was possible that DEG7 was also regulated by seed ageing. DEG10 encodes a BURP domain-containing protein, a newly identified protein that is unique to plants and plays an important role in plant abiotic stresses, development and metabolism via regulating the level of diverse proteins [32, 33]. DEG12 encodes a phenylalanine ammonia lyase homolog1 (PAL1), PAL genes was been reported involving in multiple biology process including response to environmental stress [35]. Based on the annotation of above genes, it was possible that those genes may function in ageing induced defense response, energy metabolism and the DNA/RNA and protein repair systems (Table 1).

Alternative splicing is a process whereby multiple functionally distinct transcripts are encoded from a single gene by the selective removal or retention of exons and/or introns from the maturing RNA [36, 37], which is common in many eukaryote lineages, including metazoans, fungi, plants and showing over 95% of multi-exon genes in human genome produce at least one alternatively spliced isoform [38–42]. In this study we identified 7 AS-DEGs in I178 and X178, DEG14 encodes a HSP20-like chaperones superfamily protein. DEG15 encodes a NADH dehydrogenase subunit 4, an important enzyme in the respiratory chain of all organisms having an aerobic or anaerobic electron-transport system in mitochondria [43]. DEG16 encodes an embryo defective 3012, was down-regulated and affected by ageing. DEG17 is an auxin transport protein (BIG) that in charge of the auxin polar transportation and distribution, gibberellin status in seed [44, 45]. DEG18 encodes a sucrose synthase 3, which was participate in respiration and related to plant growth [46]. DEG19 encodes a 27-kDa zein protein (zp27), specifically expressed in maize and may functions like a protease inhibitor [47]. Up-regulation of DEG14, 15, 17, 18 and 19 indicated those genes were potential ageing related genes that involved in stress response, energy metabolism, development regulation etc.

## Acknowledgements

We acknowledge the financial support from National Key R&D Program of China (2017YFD0102001-3 & 2018YFD0100900-3), National Natural Science Foundation of China (31701437 & 31771891) and the China Agriculture Research System (CARS-02-10).

## Supporting Information

**Fig S1. Data quality demonstration of RNA-Seq. A).** FPKM distribution of 2 replications of I178 and X178. **B).** Pearson correlation analysis between samples of I178 and X178.

**Fig S2. GO enrichment analysis of the DEGs.** Biology process, cellular component and the molecular function of 286 I178 DEGs **(A)** and 220 X178 DEGs **(B); C).** Only 2 co-up regulated genes detected in both I178 and X178, one of gene encodes a TBP associated factor; **D).** Total 98 common DEGs identified in both I178 and X178; **E).** 86 I178 and X178 commonly down regulated genes identified in this study, most of the genes involving in carbohydrate catabolic process and carbohydrate derivative binding.

**Fig S3. AS, ASG and AS-DEGs identified in 178. A).** 12 types of AS identified in I178 and X178 after 0d-AA and 5d-AA. **B).** Transcript isoforms (and the covered genes in brackets) that occurred AS in I178 and X178 after0d-AA and 5d-AA, the red colored are genes specifically spliced in two 178 after 5d-AA. **C).** Alternative spliced DEGs in I178 and X178 after 5d-AA. Six and one AS-DEGs were identified specifically in X178 and I178, respectively.

**Table S1. Number of reads sequenced and mapped to the maize genome.**

**Table S2. The expression of common 98 DEGs and the related biology process classification.**

**Table S3. Alternative splicing events and the genes involved in I178 and X178.**

## Author Contributions

Conceptualization: Li Li

Data curation: Li Li.

Formal analysis: Li Li, Li Xuhui.

Funding acquisition: Wang Guoying and Li Li.

Investigation: Li Li, Wangfeng and Peng Yixuan

Methodology: Li Li, Wang Feng and Peng Yixuan.

Project administration: Li Li.

Resources: Wang Guoying, Wang Jianhua and Zhang Hongwei.

Software: Li Li, Li Xuhui

Supervision: Gu Riliang and Wang Jianhua.

Validation: Li Li.

Visualization: Li Li.

Writing-original draft: Li Li.

Writing-review & editing: Li Li and Gu Riliang

## References

1. Agacka-Moldoch M, Nagel M, Doroszewska T, Lewis RS, Börne A. Mapping quantitative trait loci determining seed longevity in tobacco (Nicotiana tabacum L.). Euphytica. 2015; 202(3): 479–486.

2. Walters C. Understanding the mechanisms and kinetics of seed ageing. Seed Sci. Res. 1998; 8: 223–244.

3. Groot S, Surki A, Vos R and Kodde J. Seed storage at elevated partial pressure of oxygen, a fast method for analysing seed ageing under dry conditions. Ann. Bot. 2012; 110: 1149–1159.

4. Xin X, Lin X, Zhou Y, Chen X, Liu X, Lu X. Proteome analysis of maize seeds: the effect of artificial ageing. Physiol. Plantarum. 2011; 143, 126–138.

5. Xin X, Tian Q, Yin G, Chen X, Zhang J, Ng S. Reduced mitochondrial and ascorbate-glutathione activity after artificial ageing in soybean seed. J. Plant Physiol. 2014; 171, 140–147.

6. Nguyen TP, Cueff G, Hegedus DD, Rajjou L, Bentsink L. A role for seed storage proteins in Arabidopsis seed longevity. J. Exp. Bot. 2015; 66, 6399–6413.

7. Yin G, Xin X, Fu S, An M, Wu S, Chen X, Zhang J, He J, Whelan J, Lu X. Proteomic and carbonylation profile analysis at the critical node of seed ageing in oryza sativa. Sci. Rep. 2017; 7, 1–12.

8. He Y, Cheng J, Li X. Acquisition of desiccation tolerance during seed development is associated with oxidative processes in rice. Botanique. 2015; 94(2): 91–101.

9. Debeaujon I, Leon-Kloosterziel KM, Koornneef M. Influence of the testa on seed dormancy, germination, and longevity in Arabidopsis. Plant Physiol. 2000; 122(2): 403–414

10. Murthy UMN, Kumar PP, Sun WQ. Mechanisms of seed ageing under different storage conditions for Vigna radiata (L.) Wilczek: lipid peroxidation, sugar hydrolysis, Maillard reactions and their relationship to glass state transition. J Exp Bot. 2003; 54(384): 1057–1067.

11. Zhan J, Li W, He HY, Li CZ, He LF. Mitochondrial alterations during Al-induced PCD in peanut root tips. Plant Physiol Biochem. 2014; 75:105–113.

12. Yin G, Whelan J, Wu S, Zhou J, Chen B, Chen X, Zhang J, He J, Xin X, Lu X. Comprehensive mitochondrial metabolic shift during the critical node of seed ageing in rice. PLoS One. 2016; 11, e0148013.

13. Sano N, Kim JS, Onda Y, Nomura T, Mochida K. Okamoto M, Seo M. RNA-Seq using bulked recombinant inbred line populations uncovers the importance of brassinosteroid for seed longevity after priming treatments. Scientific report. 2017; 7: 8095 https://doi:10.1038/s41598-017-08116-5

14. Lv Y, Zhang S, Wang J, Hu Y. Quantitative Proteomic Analysis of Wheat Seeds during Artificial Ageing and Priming Using the Isobaric Tandem Mass Tag Labeling. PLOS ONE. 2016; DOI:10.1371/journal.pone.0162851 September 15, 2016

15. Wang W, Liu S, Song S, Møller IM. Proteomics of seed development, desiccation tolerance, germination and vigor. Plant Physiol. Biochem. 2015; 86, 1–15.

16. Chen X, Yin G, Börnerb A, Xin X, He J, Nage M, Liu X, Lu X. Comparative physiology and proteomics of two wheat genotypes differing in seed storage tolerance. Plant Physiology and Biochemistry. 2018; 130:455–463. https://doi.org/10.1016/j.plaphy.2018.07.022

17. Mao X, Cai T, Olyarchuk JG, Wei L. Automated genome annotation and pathway identification using the KEGG Orthology (KO) as a controlled vocabulary. Bioinformatics. 2005; 21, 3787±93. https://doi.org/10.1093/bioinformatics/bti430 PMID: 15817693.

18. Rajandeep S, Sekhon RB, Candice N, Hirsch CL, Natalia L, Shawn M, et al. Maize gene atlas developed by RNA sequencing and comparative evaluation of transcriptomes based on RNA sequencing and microarrays. PLoS ONE. 2013; 8:E61005. https://doi.org/10.1371/journal.pone.0061005. PMID:23637782.

19. Young MD, Wakefield MJ, Smyth GK, Oshlack A. Gene ontology analysis for RNA-seq: Accounting for selection bias. Genome Biol. 2010; 11, R14. https://doi.org/10.1186/gb-2010-11-2-r14 PMID: 20132535.

20. Liu Y, Zhang H, Li X, Wang F, Lyle D, Sun L, Wang G, Wang J, Li L, Gu R. Quantitative trait locus mapping for seed artificial aging traits using an F_2:3_ population and a recombinant inbred line population crossed from two highly related maize inbreds. Plant Breeding. 2018; (138):29–37. https://doi.org/10.1111/pbr.12663.

21. Shewry PR, Halford NG. Cereal seed storage proteins: Structures, properties and role in grain utilization. J Exp Bot. 2002; 53:947–958.

22. Rajjou L, Miche L, Huguet R, Job C, Job D. The use of proteome and transcriptome profiling in the understanding of seed germination and identification of intrinsic markers determining seed quality, germination efficiency and early seedling vigour. In: Navie SC, Adkins SW, Ashmore S, eds. Seeds: Biology, Development and Ecology. Oxfordshire, CAB International, 2007; 149–158.

23. Rajjou J, Lovigny Y, et al. Proteome-Wide Characterization of Seed Aging in Arabidopsis: A Comparison between Artificial and Natural Aging Protocols. Plant Physiology. 2008; 148: 620–641. https://doi.org/10.1104/pp.108.123141

24. Bewley JD. Seed germination and dormancy. Plant Cell. 1997; 9: 1055–1066.

25. Sano N, Rajjou L, North HM, Debeaujon I, Marion-Poll A, Seo M. Staying Alive: Molecular Aspects of Seed Longevity. Plant Cell Physiol. 2016; 57(4):660–74. https://doi:10.1093/pcp/pcv186.

26. Petla BP, Kamble NU, Kumar M, Verma P, Ghosh S, Singh A, Rao V, Salvi P, Kaur H, Saxena SC, Majee M. Rice PROTEIN l-ISOASPARTYL METHYLTRANSFERASE isoforms differentially accumulate during seed maturation to restrict deleterious isoAsp and reactive oxygen species accumulation and are implicated in seed vigor and longevity. New Phytol. 2016; 211(2):627–45. https://doi.org/10.1111/nph.13923.

27. Ogunola OF, Hawkins LK, Mylroie E, Kolomiets MV, Borrego E, Tang JD, Williams WP, Warburton ML. Characterization of the maize lipoxygenase gene family in relation to aflatoxin accumulation resistance. PLoS One. 2017; 12(7):e0181265. https://doi:10.1371/journal.pone.0181265. eCollection 2017.

28. Li Z, Gao Y, Lin C, Pan R, Ma W, Zheng Y, Guan Y, Hu J. Suppression of LOX activity enhanced seed vigour and longevity of tobacco (Nicotiana tabacum L.) seeds during storage. Conserv Physiol. 2018; 6(00): coy047; https://doi.org/10.1093/conphys/coy047.

29. Mochizuki S, Sugimoto K, Koeduka T, Matsui K. Arabidopsis lipoxygenase 2 is essential for formation of green leaf volatiles and five-carbon volatiles. FEBS Lett. 2016; 590(7):1017–27. https://doi:10.1002/1873-3468.12133.

30. Charton L, Plett A, Linka N. Plant peroxisomal solute transporter proteins. J Integr Plant Biol. 2019; Accepted. https://doi.org/10.1111/jipb.12790

31. Yashwanti Mudgil, Shin-Han Shiu, Sophia L. Stone, Jennifer N. Salt, and Daphne R. Goring. A Large Complement of the Predicted Arabidopsis ARM Repeat Proteins Are Members of the U-Box E3 Ubiquitin Ligase Family Plant Physiol. 2004; (134):59–66

32. Shen W, Yao X, Ye T, Ma S, Liu X, Yin X, Wu Y. Arabidopsis Aspartic Protease ASPG1 Affects Seed Dormancy, Seed Longevity and Seed Germination. Plant Cell Physiol. 2018; 59(7):1415–1431. https://doi:10.1093/pcp/pcy070.

33. Dinh SN and Kang H. An endoplasmic reticulum-localized Coffea arabica BURP domain-containing protein affects the response of transgenic Arabidopsis plants to diverse abiotic stresses. Plant Cell Rep. 2017; 36(11):1829–1839. https://doi:10.1007/s00299-017-2197-x.

34. Li Y, Chen X, Chen Z, Cai R, Zhang H, Xiang Y. Identification and Expression Analysis of BURP Domain-Containing Genes in Medicago truncatula. Front Plant Sci. 2016; 7:485. https://doi:10.3389/fpls.2016.00485. eCollection 2016.

35. Huang J, Gu M, Lai Z, Fan B, Shi K, Zhou YH, Yu JQ, Chen Z. Functional analysis of the Arabidopsis PAL gene family in plant growth, development, and response to environmental stress. Plant Physiol. 2010; 153(4):1526–38. https://doi:10.1104/pp.110.157370.

36. Chow LT, Gelinas RE, Broker TR, Roberts RJ. An amazing sequence arrangement at the 5’ends of adenovirus 2 messenger RNA. Cell. 1977; 12, 1–8. https://doi:10.1016/0092-8674(77)90180-5

37. Bush SJ, Chen L, Tovar-Corona JM, Urrutia AO. Alternative splicing and the evolution of phenotypic novelty. Philos Trans R Soc Lond B Biol Sci. 2017; 372(1713): 20150474. https://doi:10.1098/rstb.2015.0474

38. Kim N, Alekseyenko AV, Roy M, Lee C. The ASAP II database: analysis and comparative genomics of alternative splicing in 15 animal species. Nucleic Acids Res. 2007; 35, D93–D98. https://doi:10.1093/nar/gkl884

39. Grutzmann K, Szafranski K, Pohl M, Voigt K, Petzold A, Schuster S. Fungal alternative splicing is associated with multicellular complexity and virulence: a genome-wide multi-species study. DNA Res. 2014; 21, 27–39. https://doi:10.1093/dnares/dst038

40. Zhang C, Yang H, Yang H. Evolutionary character of alternative splicing in plants. Bioinform. Biol. Insights. 2016; 9(Suppl 1), 47–52. https://doi:10.4137/BBI.S33716. eCollection 2015.

41. Pan Q, Shai O, Lee LJ, Frey BJ, Blencowe BJ. Deep surveying of alternative splicing complexity in the human transcriptome by high-throughput sequencing. Nat. Genet. 2008; 40, 1413–1415. https://doi:10.1038/ng.259

42. Wang ET, Sandberg R, Luo SJ, Khrebtukova I, Zhang L, Mayr C, Kingsmore SF, Schroth GP, Burge CB. Alternative isoform regulation in human tissue transcriptomes. Nature. 2008; 456, 470–476. https://doi:10.1038/nature07509

43. Matsushita K, Otofuji A, Iwahashi M, Toyama H, Adachi O. NADH dehydrogenase of Corynebacterium glutamicum. Purification of an NADH dehydrogenase II homolog able to oxidize NADPH. FEMS Microbiol Lett. 2001; 204 (2):271–6. https://doi.org/10.1111/j.1574-6968.2001.tb10896.x. PMID: 11731134

44. Gil P, Dewey E, Friml J, Zhao Y, Snowden KC, Putterill J, Palme K, Estelle M, Chory J. BIG: a calossin-like protein required for polar auxin transport in Arabidopsis. Genes Dev. 2001; 15(15):1985–97.

45. Desgagné-Penix I, Eakanunkul S, Coles JP, Phillips AL, Hedden P, Sponsel VM. The auxin transport inhibitor response 3 (tir3) allele of BIG and auxin transport inhibitors affect the gibberellin status of Arabidopsis. Plant J. 2005; 41(2):231–42.

46. Daloso DM, Williams TC, Antunes WC, Pinheiro DP, Müller C, Loureiro ME, Fernie AR. Guard cell-specific upregulation of sucrose synthase 3 reveals that the role of sucrose in stomatal function is primarily energetic. New Phytol. 2016; 209(4):1470–83. https://doi:10.1111/nph.13704.

47. Krishnan HB, Jang S, Kim WS, Kerley MS, Oliver MJ, Trick HN. Biofortification of soybean meal: immunological properties of the 27 kDa γ-zein. J Agric Food Chem. 2011; 59(4):1223–8. https://doi:10.1021/jf103613s.

48. International rules for seed test. International seed testing association (ISTA). Switzerland: Zurich. 2018; Chapter 15–6.

49. Oliveros JC. VENNY. An interactive tool for comparing lists with Venn Diagrams. 2007; http://bioinfogp.cnb.csic.es/tools/venny/index.html.

50. Tian T, Liu Y, Yan H, You Q, Yi X, Du Z, Xu W, Su Z. AgriGO v2.0: a GO analysis toolkit for the agricultural community. Nucleic Acids Res. 2017; 45(Web Server issue): W122–W129. https://doi:10.1093/nar/gkx382.

